# Shining stars: Transgenesis and efficient metamorphosis in the sea star *Patiria miniata*

**DOI:** 10.64898/2026.08.05.742869

**Authors:** Beverly Naigles, Brian S. McGonagle, S. Zachary Swartz

**Affiliations:** Eugene Bell Center, Marine Biological Laboratory, Woods Hole, MA, USA

## Abstract

The sea star *Patiria miniata* is a widely used and powerful model organism for cell, developmental, and reproductive biology, but it has lacked genetic tools for expressing transgenes or endogenously tagging proteins. We developed a protocol to endogenously tag a broadly expressed actin gene and to express additional fluorescent markers from the same locus, using CRISPR/Cas9 genome editing. We also identified and isolated a promoter sequence of this actin gene which drives expression of transgenes. This promoter and transgene cassette can be introduced via a plasmid into the genome and persist through metamorphosis into the juvenile stage. Robust methods to induce metamorphosis that result in healthy juveniles are essential for developing stable transgenic lines and have been lacking in the field. Here we report a fast and efficient approach to induce the metamorphosis of larvae into healthy juveniles by introducing surf clam shells. Thus, we present a reliable method to generate both CRISPR/Cas9 knock-in and plasmid-integrated transgenic juvenile *P. miniata*, enabling future research on their fascinating biology, including regeneration, oogonial stem cells, metamorphosis, and more.

## Introduction

Echinoderms such as sea urchins and sea stars have long been studied as model systems for cell, developmental, and many other fields of biology, as they produce millions of accessible and optically transparent eggs and embryos (Meyer and Hinman, 2022). Studies in these animals have uncovered fundamental mechanisms of gene regulatory networks (Davidson et al., 2002a; Oliveri et al., 2002), fertilization (Giusti et al., 2003; Wessel et al., 2021), early development (Barone et al., 2022; Henson et al., 2021), embryo axis specification (McCauley et al., 2015; Sun et al., 2021; Swartz et al., 2021), organogenesis (Perillo et al., 2023), regeneration, cell biology (Bischof et al., 2017; Pal et al., 2020), and biophysics (Tan et al., 2022). Ovaries from the sea star *Patiria miniata* can be readily biopsied and cultured *ex vivo*, allowing access to oogenesis and ovarian biology, which is a unique advantage among echinoderms (Kishimoto, 2018; Paganos et al., 2026; Swartz et al., 2021). Coupled to their continuous egg production as adults, this makes them a superb system for studying reproductive biology.

Many molecular tools have been developed for sea urchins, including microinjection (Stepicheva and Song, 2014), morpholino knockdown (Meyer and Hinman, 2022), and recently, transgenesis (Jackson et al., 2024) and CRISPR/Cas9 knock-in (Lee et al., 2026; Oulhen et al., 2023). As with sea urchins, *P. miniata* has a well-annotated genome, techniques for microinjection, delivery of cargo into oocytes using VitelloTag, and both constitutive and inducible CRISPR/Cas9-mediated knockouts (Arshinoff et al., 2022; Clarke et al., 2024; Hinman and Davidson, 2007; Perillo et al., 2022; Perillo et al., 2023; Zueva and Hinman, 2023). However, prior to this work, *P. miniata* lacked tools for stable, germ line transgenesis. Thus, fluorescent markers could only be visualized for the few weeks that a microinjected mRNA persists. This meant that live investigation of late larvae, metamorphosis, and adult tissues, as well as the endogenous localization and behavior of specific proteins, had been inaccessible.

One challenge with creating transgenic *P. miniata* lines is closing the lifecycle in the laboratory with high rates of survival. It was recently shown that *P. miniata* can be cultured through its lifecycle, reaching sexual maturity in around two years (Barone et al., 2025). Previous strategies relied on retinoic acid stimulation of larvae to induce settlement and metamorphosis (Barone et al., 2025; Yamakawa et al., 2018). In our hands, however, this method results in developmental abnormalities, possibly because it bypasses an unknown natural metamorphosis trigger and mis-activates downstream developmental pathways. These findings point to the need for high-efficiency, low-toxicity approaches.

One strategy to create transgenic animals uses transposase-mediated insertion, where a transposase mediates the semi-random insertion of a sequence on an insertion plasmid into the genome (Morris et al., 2016). In the sea urchin *Lytechinus pictus*, the Minos/mariner transposase system generates stable integration in juveniles at ∼47% efficiency (Jackson et al., 2024). In other animals, a transposase is not required. For instance, *Hydra vulgaris* can stably integrate a GFP-expressing plasmid at 10-20% efficiency even in the absence of a transposase (Juliano et al., 2014; Wittlieb et al., 2006). Early work in sea urchins found that injecting linear plasmid DNA into unfertilized eggs generated an average of 56% of five-week larvae and 4-16% of post-metamorphosis juveniles that retained the exogenous DNA (Flytzanis et al., 1985; Hough-Evans et al., 1988; McMahon et al., 1985). An alternative is to microinject bacterial artificial chromosomes (BACs) that contain cis regulatory sequences driving fluorescent proteins. It is presumed that these BAC sequences concatenate and then integrate into the genome, though they may become silenced in sea urchins and be lost at metamorphosis (Davidson et al., 2002b). These approaches all require known strong promoters or large, recombineered BACs. A small fragment of the *P. miniata* Otx promoter was previously reported to express broadly in embryogenesis, but it is not clear if its expression is ubiquitous or persists past metamorphosis (Hinman et al., 2007). Another strategy uses CRISPR/Cas9-mediated knock-ins, where an endogenous protein is tagged or a transgene is inserted site-specifically into a safe harbor locus (Dickinson et al., 2013; Hruscha et al., 2013; Platt et al., 2014). Knock-ins enable visualization of endogenous proteins and ensure that proteins are expressed under endogenous regulatory control. They also do not require prior knowledge of cis-regulatory sequences.

Here, we produced *P. miniata* lines with fluorescent tags or reporters for actin, cell membranes, and histone H2B. We endogenously GFP-tagged a cytoplasmic actin gene, and then used a translational skip strategy to drive expression of several constructs from actin’s endogenous regulatory sequences at the actin locus. We then cloned a portion of this actin promoter and found that it drives ubiquitous expression of transgenes that persists past metamorphosis. Finally, we demonstrate a fast, high-efficiency, low-toxicity method of inducing metamorphosis using surf clam shells to produce transgenic juvenile *P. miniata*. These new techniques will enable visualization of organ development, metamorphosis, regeneration, and many other processes in these animals in the late larval, juvenile, and adult stages.

## Results and Discussion

### Endogenous tagging of an actin locus in *P. miniata* using CRISPR/Cas9

We first sought to endogenously tag a *P. miniata* actin gene, since we were interested in visualizing cytoskeletal behavior in adult cells. As *P. miniata* has many predicted actin genes, we analyzed our recently published ovary scRNAseq dataset and chose the actin gene LOC119719186, which is expressed in all cell-type clusters in the ovary (Paganos et al., 2026) (Fig. S1A). We designed and tested three guide RNAs (sgRNAs) against the N terminus of this locus and chose sgRNA2 for our subsequent knock-in experiments since it had the highest cutting efficiency (Fig. S1B).

We first tested a donor plasmid to knock in GFP fused to the endogenous actin protein by a flexible linker (Sheff and Thorn, 2004) (Fig. 1A). We created plasmids with ∼1 kb and ∼140 bp homology arms and injected these into oocytes, along with Cas9 mRNA and the sgRNA, which we then matured and fertilized as previously described (Perillo et al., 2023) and screened for GFP fluorescence. We found one fluorescent larva using this approach, out of hundreds screened (Fig. 1R, S1C). We next injected a double-stranded DNA PCR product with 1 kb, 140 bp, or 40 bp homology arms as our donor, and found that ∼3% of larvae injected with 1 kb or 140 bp arm donors showed fluorescence at 3 days post fertilization (dpf), while none of the 40 bp arm donor larvae did (Fig. S1C, Fig. S2). Thus, we conclude that efficiency does not increase with arms longer than 140 bp, but arms longer than 40 bp are necessary. Adding a 5’ biotin to the PCR donor increases knock-in efficiency in other systems (Paix et al., 2023). We created biotinylated donors using a 2-step handle system (Canaj et al., 2019), and found that this increased the average knock-in efficiency with 140 bp arms to from 2.2% to 7.4%, though this increase is not statistically significant (p=.064) (Fig. S1C).

**Figure 1.**
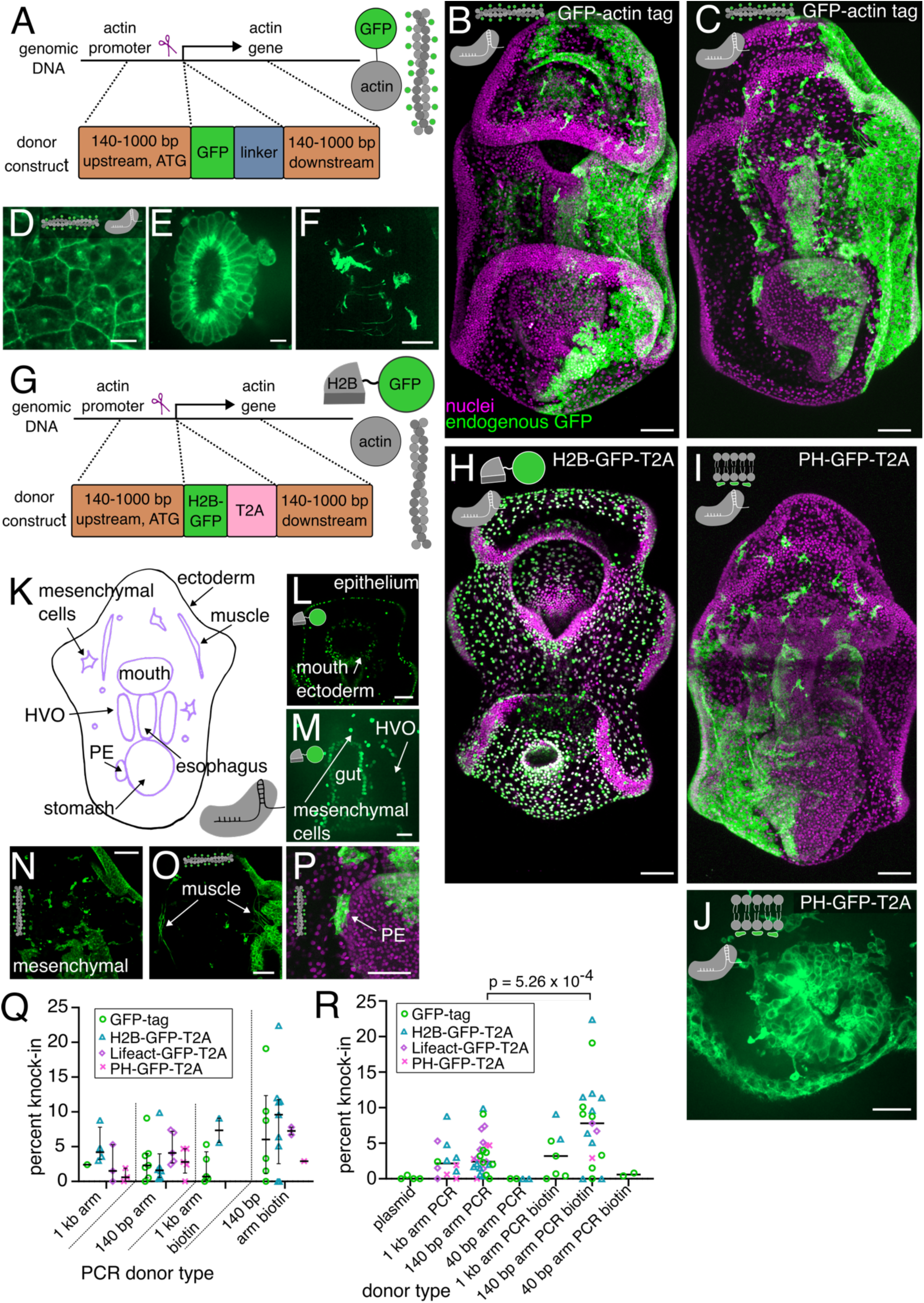
CRISPR/Cas9-mediated actin tagging and expression of exogenous markers. (**A**) Actin locus and donor construct for actin tagging. (**B-C**) Fixed 4 dpf GFP-tagged actin knock-in larvae with endogenous GFP (green) and nuclear staining (DRAQ5, magenta). (**B**) ventral view and (**C**) dorsal view of different larvae. (**D-F**) Zoom views of different cell types of 4 dpf transgenic larvae expressing endogenous GFP-tagged actin (green). (**D**) Ectodermal epithelial cells. (**E**) Intestinal epithelium. (**F**) Esophageal muscle. (**G**) Actin locus and donor construct for translational skip strategy. (**H**) Fixed 4 dpf H2B-GFP-T2A-actin knock-in larva with H2B-GFP expressed from the actin locus (green) and nuclear staining (DRAQ5, magenta), ventral view. (**I**) Fixed 4 dpf PH-GFP-T2A-actin knock-in larva with PH-GFP expressed from the actin locus (green) and nuclear staining (DRAQ5, magenta), dorsal view. (**J**) Inset of 4 dpf PH-GFP-T2A-actin knock-in larva anus showing PH localization to membranes. (**K**) Diagram of larval anatomy. (**L-P**) Zoom views of tissues in 4 dpf GFP-actin tag or H2B-GFP-T2A-actin knock-in larvae. (**L**) H2B-GFP in the epithelium and mouth ectoderm. (**M**) H2B-GFP in the gut, hydrovascular organ (HVO), and mesenchymal cells. (**N**) GFP-actin in the mesenchymal cells. (**O**) GFP-actin in the dorsal muscle (inset from C). (**P**) GFP-actin in the posterior enterocoel (PE) (inset from C) with nuclei (DRAQ5, magenta). (**Q**) Comparing each inserted sequence with different arm lengths and biotinylation states. Median and upper and lower quartiles marked. Circles indicate individual experiments. (**R**) All inserted sequences combined comparing donor arm length and biotinylation state. Circles indicate individual experiments. Bar is overall median. p-value from 2-sided t-test. Scale bar in B-C, F, H-I, L, N-P is 50 μm. Scale bar in D-E, J, M is 10 μm.

To confirm that our knock-in GFP had the expected subcellular localization for actin, we fixed larvae at 4 dpf and examined many larvae by confocal microscopy (Fig. 1B-C). We confirmed that the intracellular localization was consistent with actin and with larval phalloidin staining: localized to the periphery and surrounding the basal body of ectodermal epithelial cells, enriched at the apical side of the intestinal epithelium, and in both the esophageal and dorsal muscles (Fig. 1C-F, O, S1E). Most larvae were mosaic, implying that knock-in frequently occurred at or after the 2-cell stage. We also performed genotyping at the actin locus, which confirmed accurate insertion (Fig. S1D).

We next sought to tag histone H2B to visualize nuclei, which would be useful for live imaging experiments of cell division and migration. However, existing scRNAseq datasets did not identify a singular H2B gene which is highly expressed across all cell types, consistent with different histone genes being expressed at different stages of the lifecycle (Marzluff et al., 2006). We therefore developed a strategy to knock in H2B-GFP at the N terminus of the endogenous actin protein, separated by a T2A translational skip sequence to produce both the H2B-GFP and the endogenous actin protein as separate polypeptides (Fig. 1G, Fig. S2) (Daniels et al., 2014; Nora et al., 2017). This strategy produced larvae expressing GFP in the nucleus (Fig. 1H,M). We used a similar approach to knock in PH-GFP (Fig. 1I, J) to visualize cell membranes and Lifeact-GFP as an additional way to visualize actin (Fig. S1F-G), which both show the expected subcellular localizations. The T2A peptide therefore offers a flexible approach to drive expression of cell biological constructs.

Most individual larvae are mosaic, but we asked if this actin gene expresses in all germ layers (Fig. 1K). Using a combination of our endogenous actin tag and H2B-GFP knock-in, we collectively detected expression in the ectoderm and mouth (Fig. 1L), gut (stomach, intestines) (Fig. 1L-M), hydrovascular organ (Fig. 1M), posterior enterocoel (PE, which contains the primordial germ cells) (Fig. 1P), mesenchyme (Fig. 1N), and esophageal and dorsal muscles (Fig. 1O, F). Thus, this gene is expressed in most, if not all, major cell lineages. We then revisited our efficiency measurements with the data from these additional knocked-in sequences. We find overall that efficiency is similar between the four knocked-in sequences tested (endogenous actin tag, and the H2B-GFP, Lifeact-GFP, and PH-GFP T2A constructs), and similar between the 1 kb and 140 bp arm lengths for all donors tested (Fig. 1Q). Taking all the donor constructs together, biotinylating increases the overall efficiency with 140 bp arms from an average of 2.9% to an average of 7.8%, which when including all the data becomes statistically significant (p=.000526) (Fig. 1Q-R).

### Ubiquitous expression of transgenes by plasmid integration

Our endogenous tagging indicated that the actin gene is expressed in most or all cell types, making its promoter a strong candidate to drive a transgene cassette. We therefore created a plasmid containing 2.2 kb of sequence upstream of the 5’ UTR of the actin gene, the 1.7 kb 5’ UTR, an H2B-GFP sequence (1140 bp), and the endogenous 3’ UTR (690 bp) (Fig. 2A). We injected this plasmid into oocytes and found that 2 dpf larvae expressed H2B-GFP (Fig. 2B, Fig. S2). Most individual larvae only expressed H2B-GFP in a small number of cells at this time point, but larvae later in development frequently expressed H2B-GFP in broader domains (Fig. 2C). By inspecting multiple larvae, we determined that H2B-GFP was expressed broadly across germ layers (Fig. 2C).

**Figure 2.**
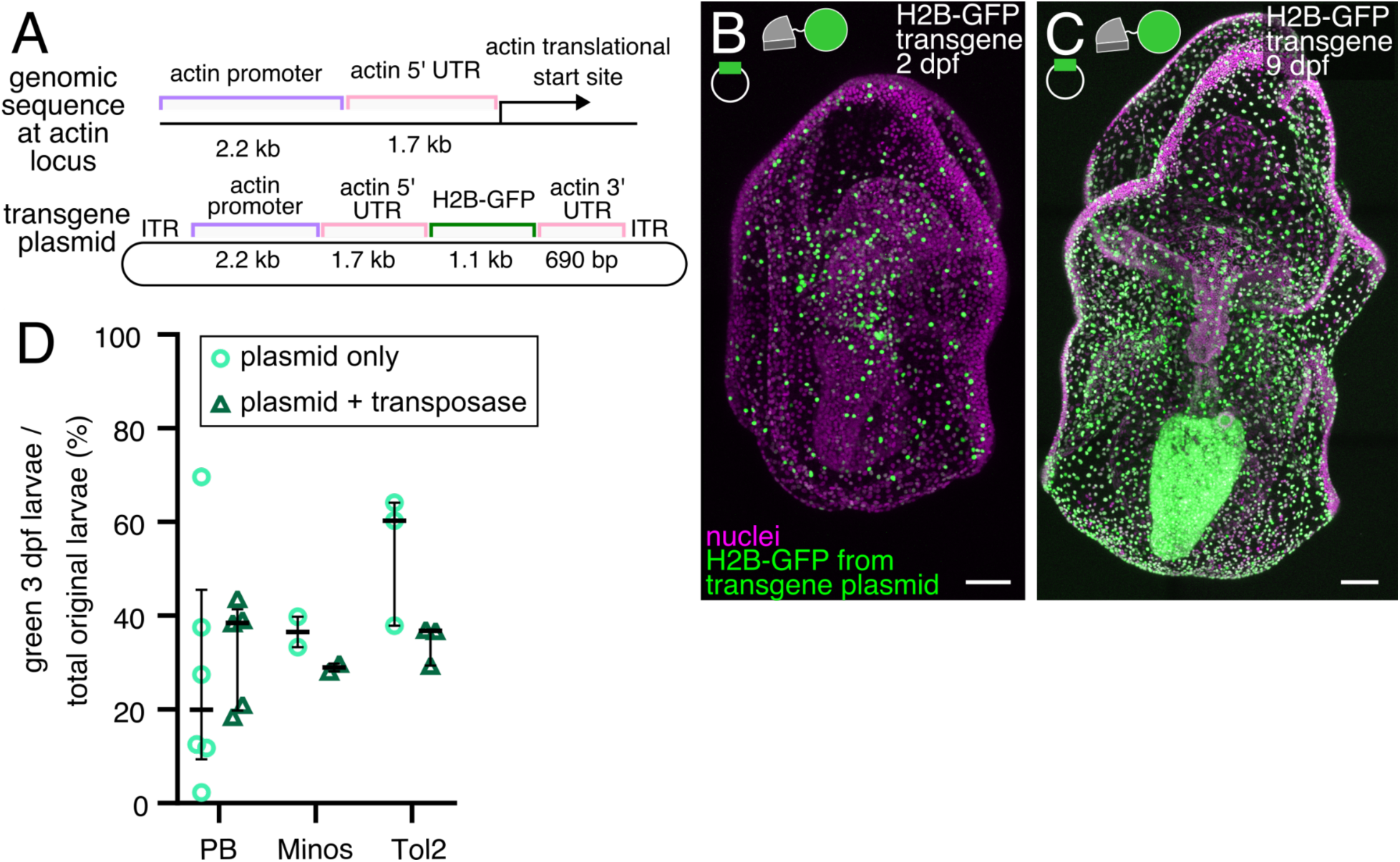
The actin promoter drives broad expression of transgenes. (**A**) Actin regulatory sequences used in the transgene plasmid. ITR is inverted terminal repeat. (**B-C**) Larvae expressing H2B-GFP transgene plasmid without transposase. Scale bar is 50 μm. (**B**) 2 dpf expressing Minos transgene plasmid. (**C**) 9 dpf expressing piggyBac transgene plasmid. (**D**) Fraction of larvae that express H2B-GFP at 3 dpf for the piggyBac (PB), Minos, and Tol2 transposase systems. Each point is a replicate, bars indicate median and upper and lower quartiles.

We next tested multiple transposase systems, including Minos, piggyBac, and Tol2, to determine which was most efficient for integration into the *P. miniata* genome (Jackson et al., 2024). We found that none of these transposases improved the expression of the H2B-GFP construct above plasmid-only controls (Fig. 2D). Thus, while this promoter holds promise for semi-random, stable integration of transgenes on plasmids, the transposase systems tested did not provide an improvement. Since some of the expression in early larvae may be from unintegrated plasmid that will be lost over time, it is necessary to track larvae to test whether expression persists through metamorphosis.

### Scallop and surf clam shells induce fast, high-efficiency, low-toxicity metamorphosis

While we were encouraged by our knock-in and plasmid-transgene-expression larvae, the efficiencies remain low and are only viable if coupled to a high-efficiency process for metamorphosis. Inspired by reports that larvae of the sea stars *Asterias forbesi* and *Asterias rubens* preferentially settle on blue mussel shells (Kalytiak-Davis and Allen, 2024), we collected a variety of local bivalve shells and presented them to competent *P. miniata* larvae. We tested shells from *Argopecten irradians* (bay scallop), *Mytilus edulis* (blue mussel), *Crepidula fornicata* (slipper limpet), *Busycon carica* (knobbed whelk), and *Spisula solidissima* (surf clam) (Fig. 3A). We found that *S. solidissima* and *A. irradians* shells robustly induced metamorphosis, at rates of 69% and 45% respectively, a significant (p=.00002 and p=.0187) increase over the no-shell control (Fig. 3B, Fig. S2).

**Figure 3.**
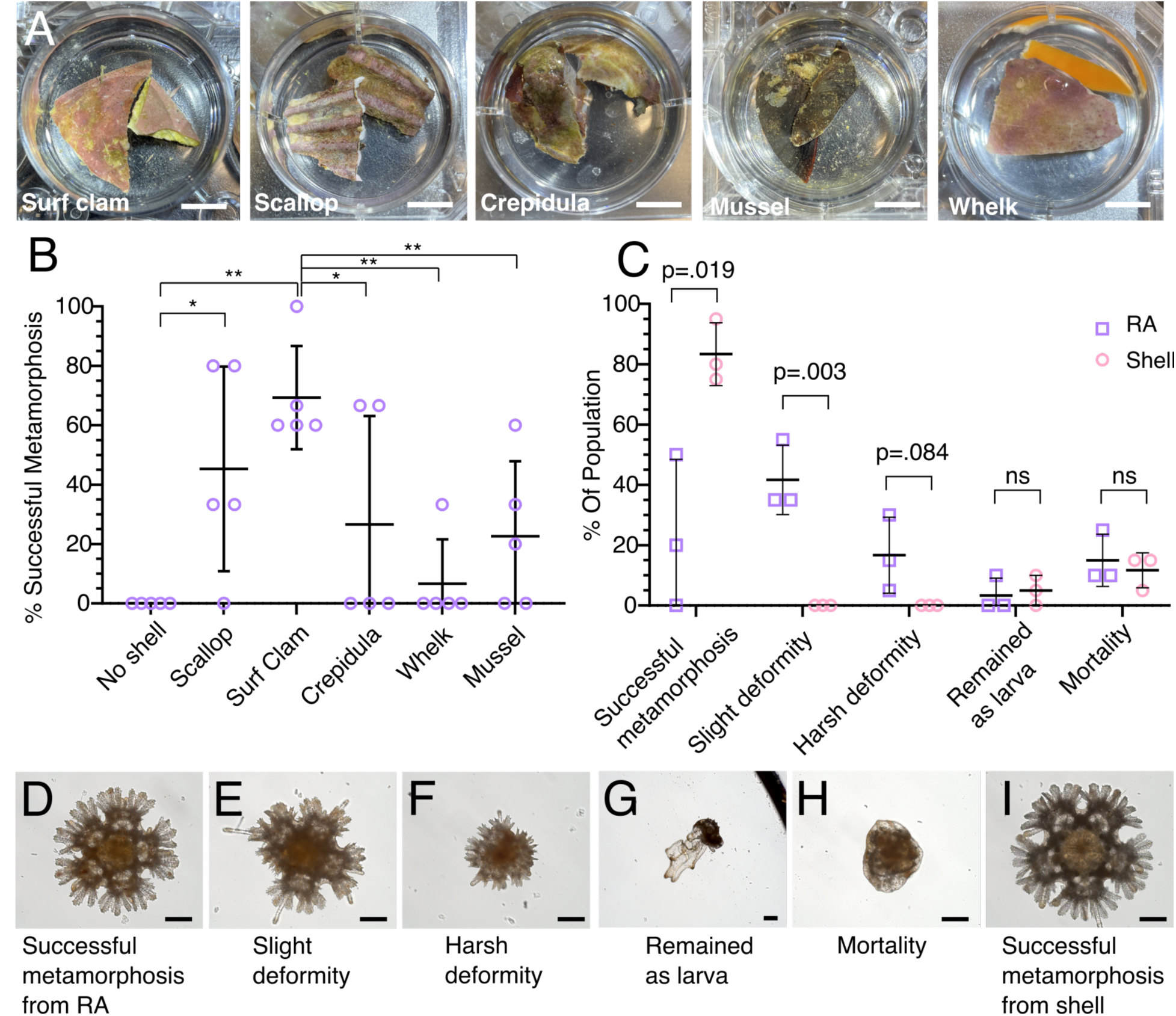
***S. solidissima* surf clam shells induce healthy metamorphosis with high efficiency**. (**A**). Example shell shards used for metamorphosis. Scale bar is 1 mm. (**B**) Fraction of larvae successfully completing metamorphosis within 1 week after exposure to each shell. Each point represents the average from 1 trial for a total of 5 trials, with 2 trials containing 3 larvae per well and 3 trials containing 5 larvae per well. * indicates p < 0.05, ** indicates p < 0.01 from a 2-tailed t-test. Relationships not marked are not statistically significant. (**C**) Fraction of original larvae showing each outcome six weeks after exposure to either RA or to a surf clam shell. Each dot represents one of three trials, each of which contained 20 competent larvae per condition. (**D-I**). Example juvenile phenotypes after metamorphosis. Scale bar is 200 μm.

Previous reports used retinoic acid (RA) to induce metamorphosis, and so we compared this to our shell approach (Barone et al., 2025; Yamakawa et al., 2018). We presented half of a competent larval cohort with shells from *S. solidissima* and exposed half to RA and scored for outcomes after six weeks. We defined slight deformities as minor issues with a single ray in the juvenile, while harsh deformities included issues with additional rays (Fig. 3D-F). Shell exposure yielded a significantly greater rate of successful metamorphosis, with no abnormal juveniles, while RA led to significant numbers of abnormal juveniles (Fig. 3C-I). A breakdown of outcomes in each trial is in Supplementary Table 1. Inducing healthy juveniles is a critical step to ensure young animals will successfully grow to reproductive adulthood. We therefore chose to induce metamorphosis with surf clam shells to rear transgenic juveniles.

### Surf clam shells lead to successful transgenic metamorphosis

We next asked whether our knock-in and transgenic larvae remain fluorescent as juveniles. After surf clam shell-induced metamorphosis, we confirmed that both knock-in actin-GFP (Fig. 4A), and H2B-GFP-T2A are expressed in juveniles, and that juveniles look healthy (Fig. 4D). We also determined that the H2B-GFP plasmid transgene remains expressed in juveniles (Fig. 4B) and that in all cases transgenic expression is above the autofluorescent background (Fig. 4C).

**Figure 4.**
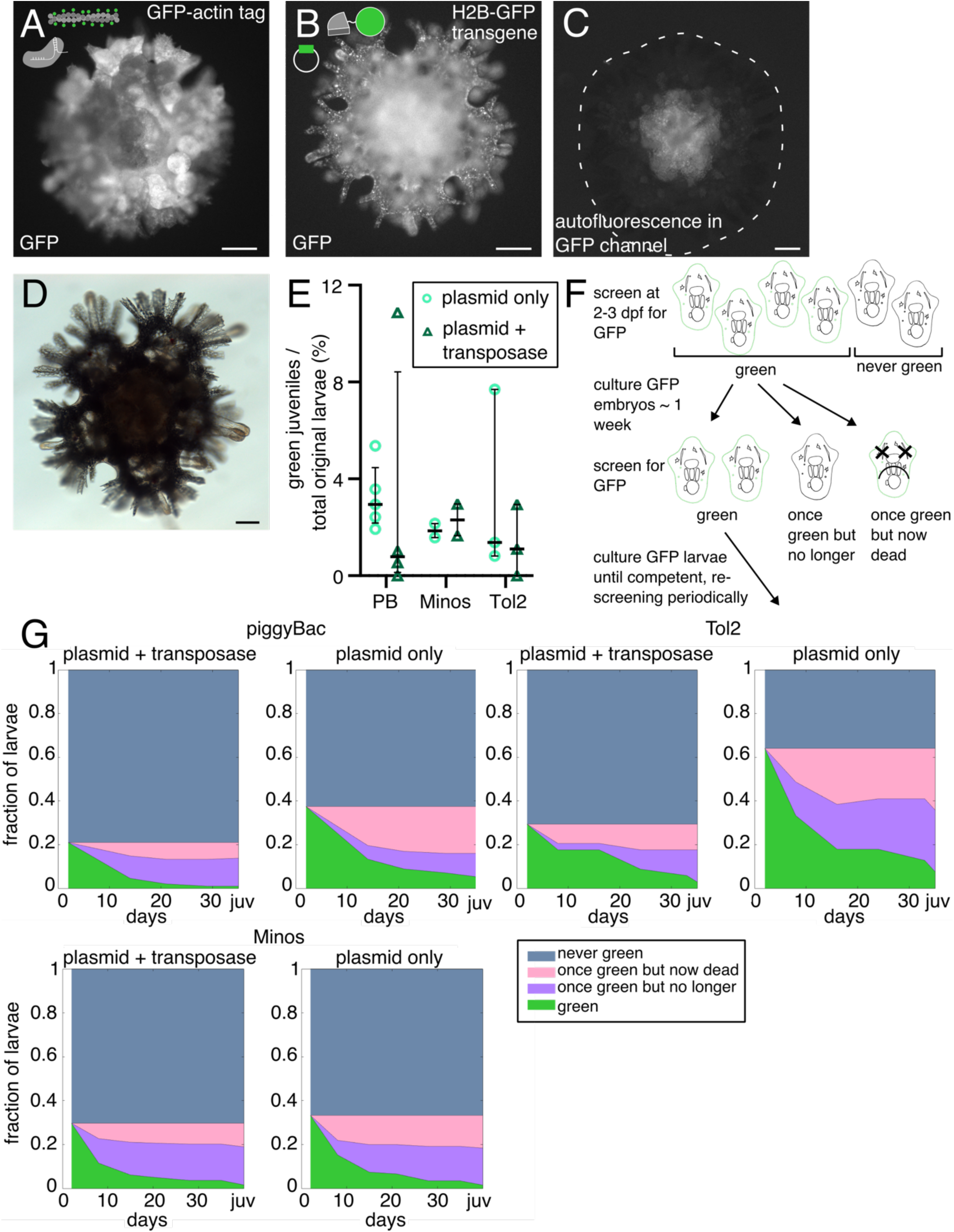
Metamorphosis of transgenic larvae produces transgenic juveniles. (**A**) GFP-actin knock-in juvenile. (**B**) H2B-GFP plasmid integration juvenile. (**C**) Non-transgenic juvenile, GFP channel. Central autofluorescent signal is the stomach, dashed white line is outline of animal. (**D**) Brightfield image of a GFP-actin tagged juvenile. (**A-D**) scale bar is 100 μm. (**E**) Fraction of juveniles that express the H2B-GFP out of all original larvae for each transposase system both with and without the transposase. Each point is one trial, sets with and without transposase were done in parallel, bars indicate median and upper and lower quartiles. (**F**) Larval screening schematic for plasmid integration experiments with and without transposase. (**G**) Fractional outcomes of all larvae injected with the H2B-GFP plasmid with and without the associated transposase. Time zero is the day of fertilization, and the juvenile timepoint is about 1 week after metamorphosis.

Expression of H2B-GFP in juveniles after injection of the plasmid with or without the transposase strongly suggests that the plasmid stably integrates. To determine if transposases improved the integration and expression of the H2B-GFP plasmids post-metamorphosis, we compared the fraction of transgenic juveniles obtained in each condition (Fig. 4E). None of the transposase systems tested increased the fraction of fluorescent juveniles compared to the plasmid alone (Fig. 4E). We also see a similar pattern of GFP loss across development in each of these conditions (Fig. 4F-G, S3A). These data are consistent with successful stable integration of the plasmid into the genome, but with no added benefit of the transposases. Thus, while future work may identify a higher-efficiency transposase system, we find that transgenic juveniles can be produced even without using a transposase.

Here we have introduced a protocol for endogenous tagging in *P. miniata* using CRISPR/Cas9, as well as for introduction of transgenes driven by the actin promoter. Going forward, these approaches could be adapted to tag additional genes, insert transgenes with tissue-specific promoters to study their spatial and temporal expression, and express cell biological markers in specific cell lineages. These approaches open new and exciting avenues of research in *P. miniata*, including questions about organogenesis, organ function, and lineage analysis in metamorphosis.

## Materials and Methods

### Research animals

Adult *Patiria miniata* were acquired from South Coast Bio (San Diego, CA) and Monterey Abalone Company (Monterey, CA) and were maintained in temperature-controlled (15°C) open-flow seawater aquaria at the Marine Resources Center at the MBL.

### Ovary and oocyte culture

Ovaries were extracted from female *P. miniata* and maintained at 15°C in 0.22 μm filtered seawater (FSW) with 50 μg/ml sulfamethoxazole and 10 μg/ml trimethoprim (ST) (diluted 1:1000 from a stock solution in DMSO). Each day ovaries were transferred to fresh FSW + ST. Oocytes were teased out from ovaries for each experiment. Sperm was extracted from male *P. miniata*, stored dry at 4°C, and used within a week.

### Oocyte microinjection and fertilization

Oocytes were microinjected with the relevant solution, incubated at 15°C in FSW + ST for 3-5 hours, matured with 1-methyladenine at a final concentration of 10 μM, and fertilized with a 1:500,000 sperm dilution. The next day, the FSW was changed. For knock-in experiments, oocytes were injected with 750 ng/uL Cas9 mRNA, 10 μM actin gRNA 2, and 30-130 ng/μL (but most commonly 60-100 ng/μL) of PCR donor, or 225 ng/μL of plasmid for the knock-in plasmid trial. For transposase experiments, oocytes were injected with 100 ng/uL plasmid + 750 ng/uL of transposase mRNA for piggyBac and 1000 ng/uL of transposase mRNA for Minos and Tol2. Transposase experiments were done by injecting in parallel the plasmid alone and the plasmid plus the transposase. Each injection experiment had 34-360 larvae per replicate.

### Larval culture

Post fertilization larvae were held in six well plates and kept at 18°C. Twice a week, wells were given water changes and fed a combination of *Isochrysis* and *Rhodomonas* sp. at a concentration of 62,500 cells/mL. Once larvae reached competency, surf clam shell shards were added to each well to induce metamorphosis.

### Larval screening

At 2 dpf (plasmid injection) or 3 dpf (knock-in), larvae were immobilized for 30 seconds in 2X seawater (1 L FSW + 30 g NaCl), transferred to regular FSW, and screened for fluorescence on a Zeiss AxioZoom V16 microscope using the GFP filter. Fluorescent larvae were sorted into a new well and cultured as described above. Fraction of green larvae was calculated as the number of green larvae divided by the total number of alive larvae.

### Guide RNA design and testing

CHOPCHOP (Labun et al., 2019) was used to find gRNAs for the N terminus of the actin locus, using settings for CRISPR/Cas9 knock-in. We selected gRNAs with high efficiencies and minimal off-target scores that were as close as possible to the ATG start site of the actin gene. These gRNAs were tested for cutting efficiency by injecting them singly into oocytes at 10 μM, along with Cas9 mRNA at 750 ng/uL and PH-GFP mRNA to identify injected embryos. The oocytes were incubated in FSW + ST for 3-5 hours, matured with 1-methyladenine at a final concentration of 10 μM, and fertilized with a 1:500,000 sperm dilution. The next day, the FSW was changed. At 3 dpf, embryos were screened for the PH-GFP and injected embryos were frozen. Genomic DNA was extracted using the QuickExtract kit (Biosearch Technologies) and the N terminal region of actin was amplified by PCR and sent for Sanger sequencing. ICE analysis was used to identify the gRNA with the highest efficiency (ICE CRISPR Analysis. 2024. EditCo Bio; [Sept 18, 2024]).

### Genotyping

Genomic DNA from larvae was extracted using the QuickExtract kit and amplified with primers that sit outside of the 140 bp arm sequence (oBN240 and oBN241) so that the primers will not bind to the injected donor DNA. The resulting product was run on a gel and imaged. To sequence these amplicons and confirm accurate insertion, the top and bottom bands were gel extracted, the PCR was repeated separately on each band to increase yield, and then the products were again gel extracted and Sanger sequenced. For the larva shown in S1D, the sequence of the insertion band was exactly as expected, and the bottom non-inserted allele was mosaic, indicating a cut that was resolved differently in different cells.

### Cloning

Genomic DNA was extracted from a *P. miniata* ovary using the QuickExtract kit (Biosearch Technologies). Knock-in donor plasmids were made with 1 kb homology arms, which were amplified from *P. miniata* genomic DNA. All primers were ordered from Genewiz (NJ, USA). Primers were designed to mutate the PAM sequence to avoid repeated Cas9 cutting. The fluorescent protein for insertion (GFP, H2B-GFP, Lifeact-GFP, PH-GFP) was amplified using PCR (NEB Q5 Master Mix, cat# M0492S) from pCS2+8 c-GFP plasmid (Gökirmak et al., 2012), pCS2+8 H2B-GFP (pZS317), pCS2+8 Lifeact-GFP (pZS229), or pCS2+8 PH-GFP (pBN034). A linker sequence of GDGAGLIN (Sheff and Thorn, 2004) or a T2A sequence from (Nora et al., 2017) was added to the DNA fragments using primer overhangs. Each PCR fragment was gel purified using a Qiagen Gel Extraction kit (cat# 28704), and then further purified using a Zymo Clean & Concentrator-5 kit (cat# D4013). The two homology arms and fluorescent protein insert were then cloned using Gibson assembly (NEB HiFi DNA Assembly Master Mix cat# E2621L) into a pUC19 backbone that had been digested with KpnI and NotI (NEB). Multiple colonies were then screened for the correct plasmid and confirmed by Sanger sequencing.

For plasmid injections, DNA was purified from bacteria using a Qiagen miniprep kit (cat# 27104) and eluted in nuclease-free water. For PCR injections, the entire cassette (left homology arm, fluorescent protein insert, right homology arm) was amplified using PCR from the 1 kb arm donor plasmid using primers to yield 1 kb arms (980 bp upstream and 987 bp downstream) (oBN037 and oBN041), 140 bp arms (135 bp upstream and 141 bp downstream) (oBN031 and oBN032), or 40 bp arms (35 bp upstream and 54 bp downstream) (oBN050 and oBN051). To 5’ biotinylate this product, primers were used to add a universal handle onto either the 1 kb arm (oBN060 and oBN061) or 140 bp arm (oBN063 and oBN063) PCR product, and then biotinylated primers (oBN056 and oBN057) against these handles were used in a PCR to produce the final biotinylated donor. For all PCR products that were injected into oocytes, DNA from the PCR was purified using the Qiagen PCR purification kit (cat# 28104), and then that product was further cleaned using a Zymo Clean & Concentrator-5 kit (cat# D4004) and eluted in nuclease-free water. As the biotinylation PCR had relatively low efficiency, 4 reactions were run and pooled together for the Qiagen PCR purification step.

To create the actin promoter plasmid, the 3.9 kb sequence upstream of the actin start codon and the annotated 690 bp 3’UTR were each amplified from the genomic DNA. H2B-GFP was amplified from pZS317. These fragments were assembled in a pUC19 vector digested KpnI/XbaI using Gibson assembly. The entire insert sequence (actin promoter, H2B-GFP, actin 3’ UTR) was then amplified with a primer to add a BamH1 site at the 5’ end so that it could be digested and ligated into transposase-specific plasmids. This amplified sequence was then digested BamHI/XbaI and ligated into Minos ITR loxp MCS lox 2272, which was a gift from Amro Hamdoun (Addgene plasmid # 218983; http://n2t.net/addgene:218983; RRID:Addgene_218983) (Jackson et al., 2024). The piggyBac ITR plasmid was created by digesting the piggyBac ITR MCS in pUC-GW-Amp plasmid (gift from Amro Hamdoun) BamHI/KpnI and ligating in the actin promoter H2B-GFP actin 3’ UTR sequence as for Minos. Ligations were done using T4 DNA ligase (NEB). The piggyBac transposase in pCS2+8 was created by amplifying the piggyBac sequence out of a T7-driven piggyBac plasmid and ligating it into pCS2+8 using AscI/AsiSI. The Tol2 ITR actin promoter H2B-GFP actin 3’ UTR plasmid was created from pBSII-SK-mTol2-MSC (gift of Karen Echeverri, originally gift from Elly Tanaka Addgene plasmid # 51817; http://n2t.net/addgene:51817; RRID:Addgene_51817) (Khattak et al., 2014) digested BamHI/ClaI. The insert sequence was PCR amplified from the Minos actin promoter H2B-GFP actin 3’ UTR plasmid using primers to add Gibson overhangs and assembled with the backbone by Gibson assembly. The Tol2 transposase in pCS2+8 was created by amplifying the Tol2 sequence from pT3TS-Tol2 (gift of Carrie Albertin, originally gift from Stephen Ekker Addgene plasmid # 31831; http://n2t.net/addgene:31831; RRID:Addgene_31831) (Balciunas et al., 2006) to add Asc1 and AsiS1 restriction sites, which was then digested and ligated into pCS2+8.

Oligos used in this study can be found in Supplementary Table 2.

### Plasmids created in this study

| Plasmid number | Name |
| --- | --- |
| pBN013 | pUC19 GFP-actin 1 kb arm donor |
| pBN027 | pUC19 H2B-GFP-T2A-actin 1 kb donor |
| pBN036 | pUC19 PH-GFP-T2A-actin 1 kb donor |
| pBN037 | pUC19 Lifeact-GFP-T2A-actin 1 kb donor |
| pBN047 | Minos ITR loxp actin prom H2B-GFP actin 3' UTR lox2272 |
| pBN050 | piggyBac ITR actin prom H2B-GFP actin 3' UTR in pUC-GW-Amp |
| pBN051 | pCS2+8 piggyBac transposase |
| pBN070 | pCS2+8 Tol2 transposase |
| pBN071 | pBSII Tol2 ITR actin prom H2B-GFP actin 3' UTR |

### Guide RNAs used in this study

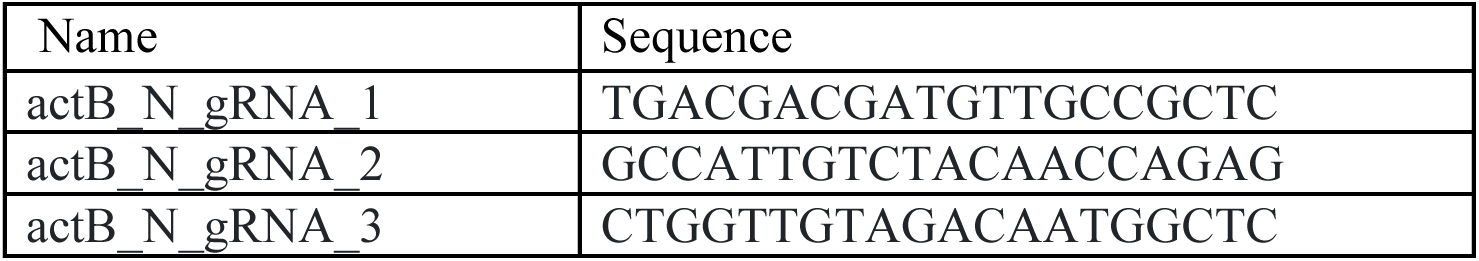

### In vitro transcription

Minos transposase mRNA was created from the pBlueSKMimRNA plasmid, which was a gift from Michalis Averof (Addgene plasmid # 102535; http://n2t.net/addgene:102535; RRID:Addgene_102535) (Pavlopoulos et al., 2004). Tol2 transposase and piggyBac transposase mRNA were created from pBN051 and pBN070. Cas9 mRNA was made from pCS2-3xFLAG-NLS-SpCas9-NLS, which was a gift from Yonglong Chen (Addgene plasmid # 51307; http://n2t.net/addgene:51307; RRID:Addgene_51307) (Guo et al., 2014). All plasmids were digested with NotI (NEB), column purified, and then in vitro transcribed using the Promega RiboMax SP6 in vitro transcription kit (Promega cat# P1280) and poly-A-tailed using the Invitrogen poly-A tailing kit (Thermo Fisher cat# AM1350).

### Fixed larval imaging

Transgenic larvae were fixed in 4% paraformaldehyde (PFA) in FSW for 20 minutes, then permeabilized in PBST (0.1% Triton) for 20 minutes, and then incubated in PBST + 1:1000 DRAQ5 (Invitrogen cat# 65-0880-92) for 30 minutes. Larvae were then imaged on a Zeiss 780 or Zeiss spinning disk confocal microscope.

### Metamorphosis on shells

Shells were collected from Vineyard Sound, MA, USA using a dredge on board the R/V Gemma. Shells were visually analyzed and shells with visible microbiome growth were selected as metamorphosis substrates. They were then stored until use in ambient sea water in a flow-through system. Shells were broken into smaller pieces and set up in 5 mL of FSW in six-well plates. Competent larvae were then added to each well. Each well was checked after one week and all larvae that successfully completed metamorphosis were counted and removed.

### Metamorphosis retinoic acid trials

*Spisula solidissima* shells were collected, selected, and stored as described above. Shells were broken into smaller pieces and added to 15 mL of FSW in 55 mm petri dishes. A 0.1 μM retinoic acid (RA) solution was made in 15 mL of FSW in small petri dishes. RA-exposed larvae were left in the solution for 48 hours and then removed and added to clean FSW (Barone et al., 2025). Shell-exposed larvae were left with shells for 7 days and any successfully metamorphosed juveniles were moved to clean FSW. Juveniles were then monitored for 6 weeks post-exposure while noting any mortalities or degradation to health, and the final outcomes were reported after 6 weeks.

### Statistics

All t-tests were two-sample, two-tailed unpaired t-tests assuming equal variances.

A more detailed protocol for creating the knock-in juveniles can be found in the Supplementary Information.

## Author contributions

Conceptualization: B.N., S.Z.S.; Methodology: B.N., B.S.M.; Formal analysis: B.N.; Investigation: B.N., B.S.M.; Writing – original draft: B.N., B.S.M.; Writing – review & editing: B.N.; B.S.M., S.Z.S.; Supervision: S.Z.S.; Project administration: S.Z.S.; Funding acquisition: B.N.,S.Z.S.

## Acknowledgements

We thank current and former Swartz lab members, including Jamie MacKinnon, Akshay Kane, and Periklis Paganos, for helpful discussions. We thank Margherita Perillo for helpful discussions, larval anatomy advice, and for not telling Beverly she was crazy when she first kicked around this idea. We thank Amro Hamdoun for helpful discussions and for sharing plasmids, Karen Echeverri for pBSII SK Tol2 MCS, and Carrie Albertin for the pT3TS-Tol2 plasmid. We thank Jan Soroczyński for manuscript feedback and pointing us to the universal biotinylated handle approach. We thank Lisa Abbo, Nolan Gibbons and the staff of the MBL Marine Resources Center for assistance with sea star husbandry and shell collection aboard the R/V Gemma. We thank the MBL Central Microscopy Facility for imaging resources. We thank the MBL Keck Facility for Sanger sequencing.

## Funding

This work was supported by a Faculty Recruitment Gift from the Hibbitt Foundation and National Institutes of Health NIGMS MIRA R35GM1624422. S. Zachary Swartz, PhD is a Pew Scholar in the Biomedical Sciences, supported by The Pew Charitable Trusts. BN was supported by National Institutes of Health NICHD F32HD117487.

## Data and Resource Availability

Plasmids and their sequences have been deposited on addgene. Transgenic sperm, embryos, and adults will be shared upon reasonable request as soon as it is experimentally feasible, depending on the growth of the lines.

## Notes

### Competing Interest Statement

The authors have declared no competing interest.

