## Supplementary Information for "Shining stars: Transgenesis and efficient metamorphosis in the sea star *Patiria miniata*"

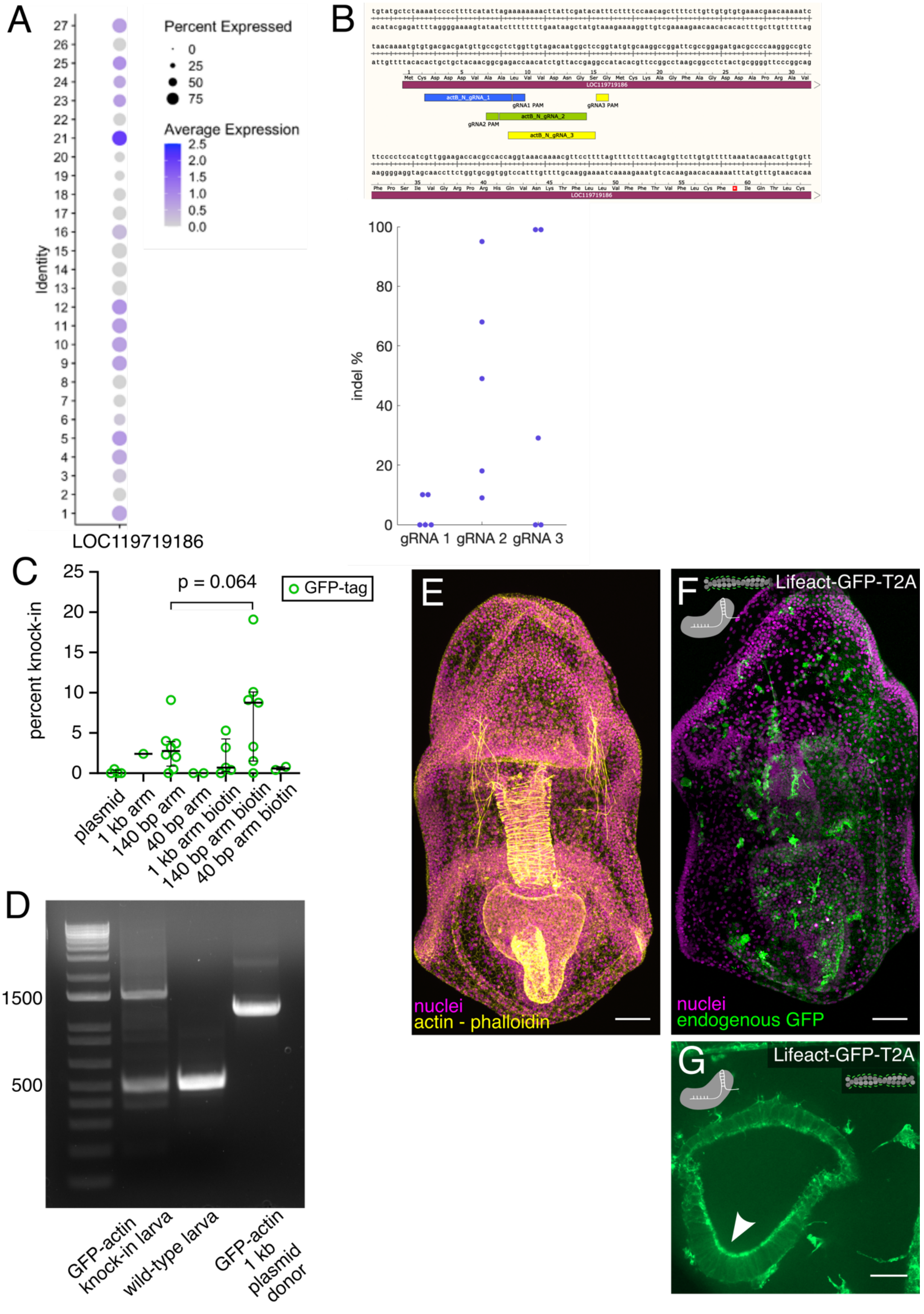
A
Percent Expressed
0
25
50
75
Average Expression
2.5
2.0
1.5
1.0
0.5
0.0
Identity
LOC119719186
B
LOC119719186
actB_N_gRNA_1
gRNA1 PAM
gRNA3 PAM
gRNA2 PAM
actB_N_gRNA_2
actB_N_gRNA_3
indel %
gRNA 1
gRNA 2
gRNA 3
C
p = 0.064
GFP-tag
percent knock-in
plasmid
1 kb arm
140 bp arm
40 bp arm
1 kb arm biotin
140 bp arm biotin
40 bp arm biotin
D
1500
500
GFP-actin knock-in larva
wild-type larva
GFP-actin 1 kb plasmid donor
E
nuclei
actin - phalloidin
F
Lifeact-GFP-T2A
nuclei
endogenous GFP
G
Lifeact-GFP-T2A

**Figure S1.** (**A**) Average expression intensity and percent of cells expressing actin LOC119719186 in the 27 cell clusters in the *P. miniata* ovary, from Paganos et al., 2026. (**B**) Snapgene diagram of the N terminus of the LOC119719186 locus with the 3 tested gRNAs indicated. Scatter plot showing the indel percentage in each of 5 larvae injected with each gRNA. (**C**) Knock-in efficiency for the GFP-actin tag donor with different arm lengths and biotinylation states. Bars indicate median and upper and lower quartiles. Circles indicate individual experiments. p-value from 2-sided t-test. (**D**) Genotyping gel showing in lane 1: knock-in and wild-type actin locus band in GFP-actin knock-in larva made using a 140 bp arm donor; in lane 2: wild-type actin locus band in wild-type larva; in lane 3: inserted sequence amplified from 1 kb arm donor plasmid as a positive control. (**E**) 4 dpf larva stained with phalloidin (yellow) to visualize actin and DRAQ5 (magenta) to visualize DNA. Scale bar is 50 µm. (**F**) Fixed 4 dpf Lifeact-GFP-T2A-actin knock-in larva with Lifeact-GFP expressed from the actin locus (green) and nuclear staining (DRAQ5, magenta). Scale bar is 50µm. (**G**) Inset of 4 dpf Lifeact-GFP-T2A-actin knock-in larva showing Lifeact localization to the apical side of the mouth epithelium. Scale bar is 10µm.

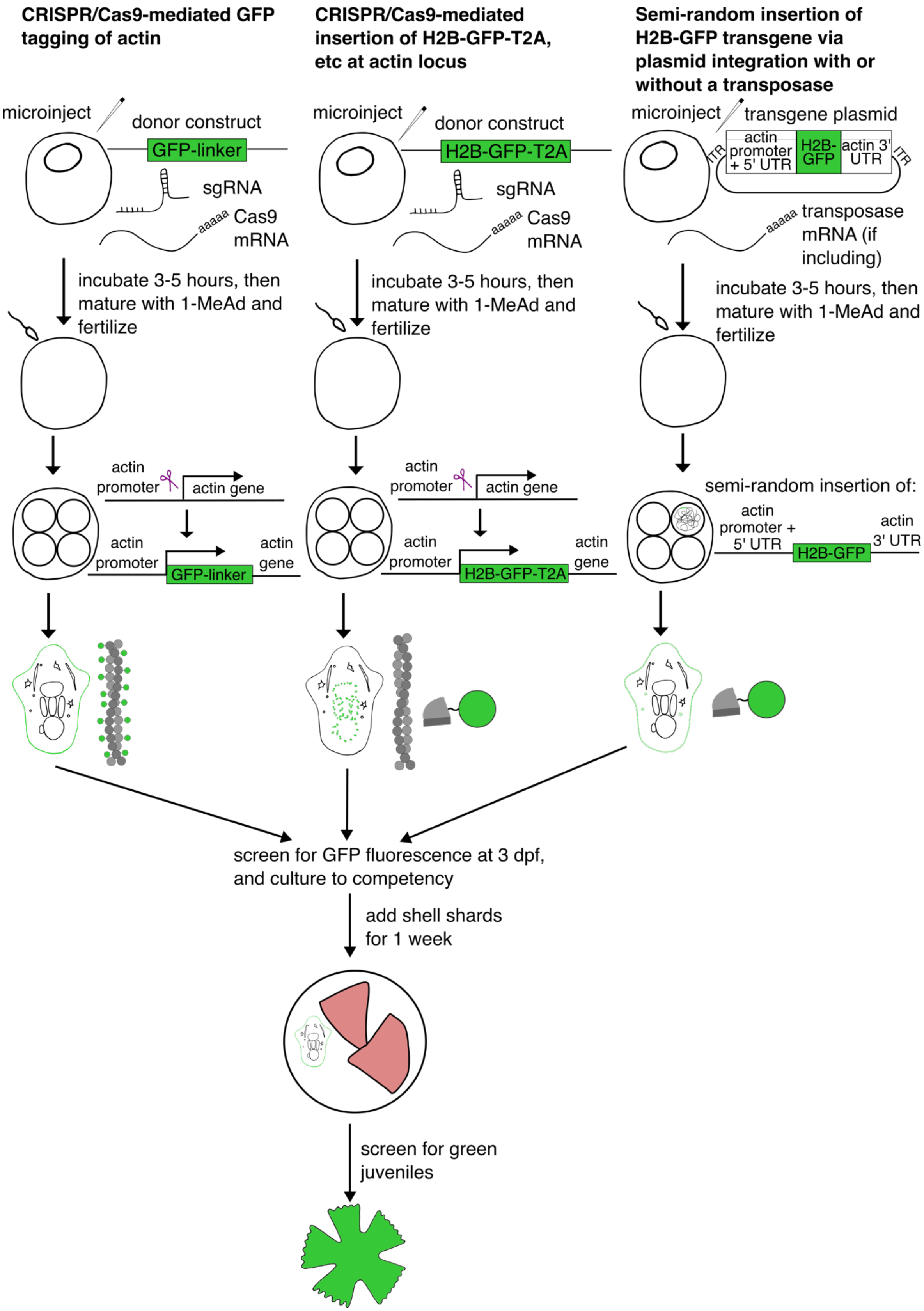


**Figure S2**. Experimental flowchart for the three types of transgenic animals created in this study.

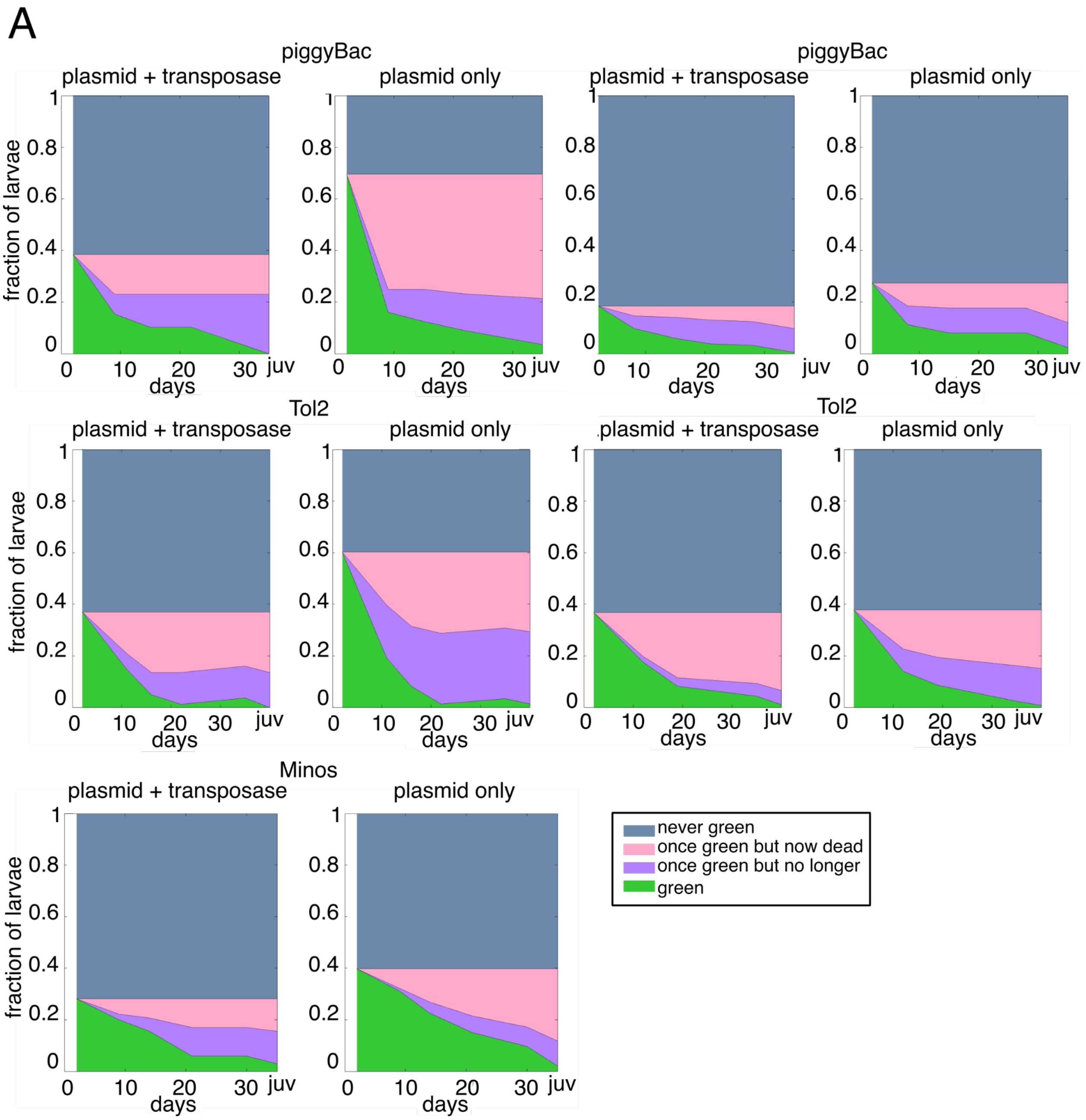


**Figure S3.** (**A**) Outcomes of all larvae from oocytes injected with the H2B-GFP plasmid with and without the associated transposase. Combined with Fig. 4G this shows all trials run. Day 0 is the day of fertilization. Juvenile timepoint is within a week of metamorphosis, which varies slightly between conditions. Each condition began with 34-355 animals.

**Supplementary Table 1 - RA vs shell experiment animal outcomes by trial**

**Percent of animals experiencing each outcome**

**RA**

| | **Trial 1** | **Trial 2** | **Trial 3** |
|---|---|---|---|
| **Successful metamorphosis** | 0 | 20 | 50 |
| **Slight deformity** | 35 | 55 | 35 |
| **Harsh deformity** | 30 | 15 | 5 |
| **Remained as larva** | 10 | 0 | 0 |
| **Mortality** | 25 | 10 | 10 |

**Shell**

| | **Trial 1** | **Trial 2** | **Trial 3** |
|---|---|---|---|
| **Successful metamorphosis** | 95 | 75 | 80 |
| **Slight deformity** | 0 | 0 | 0 |
| **Harsh deformity** | 0 | 0 | 0 |
| **Remained as larva** | 0 | 10 | 5 |
| **Mortality** | 5 | 15 | 15 |

**Supplementary Table 2 - Oligos used in this study**

| Primer name | Sequence | Purpose |
|---|---|---|
| oBN031_actin_N_cross_v2_F | GGATCGTGTTTGATGTTTGAGA | PCR across gRNA cut site for ICE analysis, also PCR to make 140 bp arm donor |
| oBN032_actin_N_cross_v2_R | AAGGAACGTTTTGTTTACCTGG | PCR across gRNA cut site for ICE analysis, also PCR to make 140 bp arm donor |
| oBN037_actin_Larm_1kb_F | gacggccagtgaattcgagctcggtaccGAATGCCAAATTGAAGTTTT | PCR 1kb left homology arm from *P. miniata* genome, also used to amplify 1 kb arm PCR donor from plasmid |
| oBN038_actin_Larm_R | CATTTTGTTAGATTTTTGTTCG | PCR 1kb left homology arm from *P. miniata* genome |
| oBN040_actin_Rarm_F | cggcctgatcaacatgtgtgacgacgatgttgctgCTCTGGTTGTAGACAATGGCTCC | PCR 1kb right homology arm from *P. miniata* genome |
| oBN041_actin_Rarm_1kb_R | gcttgcatgcctgcaggtcgactctagaCTCGGCTGAAAAGAGCAAAACAA | PCR 1kb right homology arm from P miniata genome, also used to amplify 1 kb arm PCR donor from plasmid |
| oBN043_GFP_for_actin_F | gaaacgaacaaaaatctaacaaaATGAGTAAAGGAGAAGAACTTTTC | PCR GFP to make GFP-actin tag donor |
| oBN044_GFP_for_actin_R | tcgtcacacatgttgatcaggccggcgccgtcgccTTTGTATAGTTCATCCATGCC | PCR GFP to make GFP-actin tag donor |
| oBN050_actin_40bp_F | cttgttgtgtgtgaaacgaacaa | Amplify 40 bp arm cassette off 1 kb donor plasmid |
| oBN051_actin_40bp_R | cttgcacataccggagcc | Amplify 40 bp arm cassette off 1 kb donor plasmid |
| oBN056_univ_biotin_handle_F | /5Bio/G*G*G*A*A*CCTCTTCTGTAACTCCTTAGC | Universal primer from handle to add biotin |
| oBN057_univ_biotin_handle_F | /5Bio/C*C*T*G*A*G*GGCAAACAAGTGAGCAGG | Universal primer from handle to add biotin |
| oBN060_actin_1kb_arm_handle_F | gggaacctcttctgtaactccttagcGAATGCCAAATTGAAGTTTT | Amplify 1 kb arm donor cassette from donor plasmid and add handle for future biotinylation |
| oBN061_actin_1kb_arm_handle_R | cctgagggcaaacaagtgagcaggCTCGGCTGAAAAGAGCAAAACAA | Amplify 1 kb arm donor cassette from donor plasmid and add handle for future biotinylation |

| | | |
|---|---|---|
| oBN062_actin_200bp_v2_arm_handle_F | gggaacctcttctgtaactccttagcGGATCGTGTTTGATGTTTGAGA | Amplify 140 bp arm donor cassette from donor plasmid and add handle for future biotinylation |
| oBN063_actin_200bp_v2_arm_handle_R | cctgagggcaaacaagtgagcaggAAGGAACGTTTTGTTTACCTGG | Amplify 140 bp arm donor cassette from donor plasmid and add handle for future biotinylation |
| oBN064_actin_40bp_arm_handle_F | gggaacctcttctgtaactccttagcCTTGTTGTGTGTGAAACGAACAA | Amplify 40 bp arm donor cassette from donor plasmid and add handle for future biotinylation |
| oBN065_actin_40bp_arm_handle_R | cctgagggcaaacaagtgagcaggCTTGCACATACCGGAGCC | Amplify 40 bp arm donor cassette from donor plasmid and add handle for future biotinylation |
| oBN094_H2B_for_actin_v2_F | ctaacaaaATGACGAAAGCATCCAGTCG | Amplify H2B-GFP from pZS317 |
| oBN067_GFP_to_T2A_R | gtcaccgcatgttagcagacttcctctgccctctccactgccTTTGTATAGTTCATCCATGCCATGTG | Amplify H2B-GFP from pZS317, adding a T2A sequence at the end |
| oBN068_T2A_actin_Rarm_F | agtctgctaacatgcggtgacgtcgaggagaatcctggcccaatgtgtgacgacgatgttgcTgCTCTGGTTGTAGACAATGGCTCC | Used with oBN041 to amplify the 1 kb right homology arm to add the T2A sequence |
| oBN093_actin_Larm_v2_R | ggatgctttcgtCATTTTGTTAGATTTTTGTTCG | Use with oBN037 to amplify the left homology arm to create the H2B-GFP 1 kb arm donor plasmid |
| oBN116_Lifeact_for_actin_F | ctaacaaaATGGGCGTGGCCGACTT | Use with oBN067 to amplify Lifeact-GFP from pZS229 |
| oBN117_actin_Larm_v2_Lifeact_R | gtcggccacgcccATTTTGTTAGATTTTTGTTCG | Use with oBN037 to amplify left 1 kb homology arm from *P. miniata* genomic DNA with Gibson overhang to Lifeact |
| oBN118_PH_for_actin_F | ctaacaaaATGGACTCGGGCCGGG | Use with oBN067 to amplify PH-GFP from pBN034 |
| oBN119_actin_Larm_v2_PH_R | ccggcccgagtcCATTTTGTTAGATTTTTGTTCG | Use with oBN037 to amplify left 1 kb homology arm from *P. miniata* genomic DNA with Gibson overhand to PH |
| oBN129_actin_3.9kb_promoter_v2_F | gacggccagtgaattcgagctcggtaccGTCAAAACCTTTACATGTGC | Amplify ~2kb actin promoter and ~1.7 kb actin 5' UTR from genomic DNA |
| oBN123_actin_3.7kb_promoter_R | ccacgactggatgctttcgtcatTTTGTTAGATTTTTGTTCGTTTC | Amplify ~2kb actin promoter and ~1.7 kb actin 5' UTR from genomic DNA |

| | | |
|---|---|---|
| oBN124_actin_3'UTR_F | gcatggatgaactatacaaataaACAAACTGTAAAAAACCCAAC | Amplify actin 3' UTR from genomic DNA |
| oBN125_actin_3'UTR_R | gcttgcatgcctgcaggtcgactctagaACTCTGCAAGCTTCAATAAA | Amplify actin 3' UTR from genomic DNA |
| oBN126_H2B _F | ATGACGAAAGCATCCAGTCG | Amplify H2B-GFP from pZS317 to assemble with actin promoter and 3' UTR |
| oBN127_GFP_ R | TTATTTGTATAGTTCATCCATGCCATGTG | Amplify H2B-GFP from pZS317 to assemble with actin promoter and 3' UTR |
| oBN149_actin_3.9kb_promoter_v2_BamH1_F | gacggccagtgaattcgagctcggtaccggatccGTCAAAACCTTTACATGTGC | Amplify off actin promoter - H2B-GFP - actin 3' UTR plasmid to add a 5' BamH1 site |
| oBN166_PBtransposase_F | gcgataggcgcgccATGGGTAGTTCTTTAGACGATG | Amplify PB transposase from T7 expression plasmid to ligate into SP6 expression plasmid |
| oBN167_PBtransposase_R | gcgatagcgatcgcTCAGAAACAACTTTGGCACA | Amplify PB transposase from T7 expression plasmid to ligate into SP6 expression plasmid |
| oBN208_actin_3.9kb_promoter_v2_BamH1_GibsonTol2_F | gtggcggccgctctagaactagtggatccGTCAAAACCTTTACATGTGC | Amplify actin promoter - H2B-GFP - actin 3' UTR to add Gibson overhangs into Tol2 ITR MSC plasmid |
| oBN209_actin_3'UTR_Cla1_GibsonTol2_R | ggccccccctcgaggtcgacggtatcgatACTCTGCAAGCTTCAATAAA | Amplify actin promoter - H2B-GFP - actin 3' UTR to add Gibson overhangs into Tol2 ITR MSC plasmid |
| oBN210_Tol2transposase_F | gcgataggcgcgccATGGAGGAAGTATGTGATTC | Amplify Tol2 transposase from T3 expression plasmid to ligate into SP6 expression plasmid |
| oBN211_Tol2transposase_R | gcgatagcgatcgcCTACTCAAAGTTGTAAAACCTC | Amplify Tol2 transposase from T3 expression plasmid to ligate into SP6 expression plasmid |
| oBN240_actin_N_cross_v9_F | GTAGGGCCCTTGGACAATAAG | Genotype insertion at the actin locus |
| oBN241_actin_N_cross_v9_R | AACACAAGAACACTGTAAAGAAAAC | Genotype insertion at the actin locus |

***P. miniata* CRISPR/Cas9 knock-in protocol**
**August 3, 2026**

1. Prepare the gRNAs
    a. Order gRNAs from Synthego, resuspend to 100 μM in nuclease-free water, aliquot in 5μL aliquots, store at -80°C
2. Prepare the Cas9 mRNA
    a. Miniprep Cas9 mRNA from addgene # 51307
    b. Digest this miniprep with NEB NotI for 2 hours at 37°C
    c. Run 2 μL of digest on an agarose gel to confirm linearization
    d. PCR purify the rest of the digest (we use the Qiagen PCR purification kit)
    e. Perform in vitro transcription and polyA-tailing according to kit instructions (we use the Promega RiboMAX SP6 kit # P1280 and Invitrogen polyA tailing kit #AM1350).
3. Prepare biotinylated PCR product
    a. Amplify the PCR product from the 1 kb arm donor plasmid using primers for the desired arm length to add handles for biotinylation - for 140 bp this is oBN062 and oBN063.
    b. Run 2 μL of product on a gel to confirm size, and then column purify the rest.
    c. Set up and run 4 x 50 μL PCR reactions using this purified PCR product as a template, and amplify using oBN056 and oBN057.
    d. Pool these PCR products, and run 2 μL on a gel to check sizing. Column purify the combined products using a Qiagen PCR purification kit, followed by a Zymo DNA Clean & Concentrator kit. Elute from the final column in 12 μL nuclease-free water.
4. Prepare injection mix
    a. In a 2 μL volume, combine Cas9 mRNA to a final concentration of 750 ng/μL, 0.2 μL of the 100 μM actin gRNA 2, and fill the rest of the volume with biotinylated PCR donor, aiming for a final concentration of 60-130 ng/μL of PCR donor DNA.
5. Inject oocytes
    a. Inject *P. miniata* oocytes with this mix. We aim for the germinal vesicle, though we have not determined whether injecting the GV or the cytoplasm gives better results and this does not seem to be a critical detail. Recover oocytes into FSW + ST.
    b. Incubate injected oocytes at 15°C for 3-5 hours.
    c. Mature oocytes with 10 μM 1-methyladenine and incubate about 45 minutes until the germinal vesicle is no longer visible.
    d. Fertilize oocytes with *P. miniata* sperm at a final dilution of 1:500,000.
    e. Once fertilization envelopes have risen, move fertilized oocytes to fresh FSW.
    f. The next morning, change the water of the embryos, removing any dead or unfertilized oocytes or embryos.
6. At 3 dpf, immobilize larvae by submerging them in 2X seawater (1 L FSW + 30 g NaCl) for 30 seconds and then recovering them into a FSW wash before recovering them into a new well of FSW. They will remain immobilized for 30-45 minutes. Screen for GFP

fluorescence on a microscope, we use an upright Zeiss AxioZoom V16. Separate green larvae into a new well.

7. Culture green larvae to competency, changing their water and feeding them twice a week.
8. When competent, add surf clam shell shards into a well with new FSW. Leave at room temperature for 1 week.
9. Screen juveniles for GFP fluorescence on AxioZoom V16.
